# Stiefel Manifold Dynamical Systems for Tracking Representational Drift

**DOI:** 10.64898/2026.03.07.710319

**Authors:** Hyun Dong Lee, Aditi Jha, Stephen E. Clarke, Michael P. Silvernagel, Paul Nuyujukian, Scott W. Linderman

## Abstract

Understanding neural dynamics is crucial for uncovering how the brain processes information and controls behavior. Linear dynamical systems (LDS) are widely used for modeling neural data due to their simplicity and effectiveness in capturing latent dynamics. However, LDS assumes a stable mapping from the latent states to neural activity, limiting its ability to capture representational drift—gradual changes in the brain’s representation of the external world. To address this, we introduce the Stiefel Manifold Dynamical System (SMDS), a new class of model designed to account for drift in neural representations across trials. In SMDS, emission matrices are constrained to be orthonormal and evolve smoothly over trials on the Stiefel manifold—the space of all orthonormal matrices—while the dynamics parameters are shared. This formulation allows SMDS to leverage data across trials while accounting for non-stationarity, thus capturing the underlying neural dynamics more accurately compared to an LDS. We apply SMDS to both simulated datasets and neural recordings across species. Our results consistently show that SMDS outperforms LDS in terms of log-likelihood and requires fewer latent dimensions to capture the same activity. Moreover, SMDS provides a powerful framework for quantifying and interpreting representational drift. It reveals a gradual drift over the course of minutes in the neural recordings and uncovers varying drift rates across dimensions, with slower drift in behaviorally and neurally significant dimensions.

## 1 Introduction

Latent dynamical systems are widely used in systems neuroscience to understand the evolution of neural activity over time (Smith and Brown, 2003; Eden et al., 2004; Macke et al., 2011; Archer et al., 2014; Gao et al., 2016; Linderman et al., 2017; Zhao and Park, 2017; Pandarinath et al., 2018; Duncker and Sahani, 2018; Glaser et al., 2020; Bong et al., 2020; Vyas et al., 2020; Zoltowski et al., 2020; Hu et al., 2024). A fundamental insight driving this approach is that high-dimensional neural population activity is often confined to low-dimensional manifolds (Vyas et al., 2020; Duncker and Sahani, 2021). Latent dynamical systems leverage this insight by modeling high-dimensional neural data with low-dimensional latent states, revealing underlying dynamical computations and how they result in observed behavior. Linear dynamical systems (LDS) are a common choice in this family, owing to their computational simplicity and ability to effectively approximate neural dynamics (Paninski et al., 2010; Macke et al., 2011; Archer et al., 2014; Gao et al., 2016; Soldado-Magraner et al., 2024; Jha et al., 2024).

These models, however, have a key limitation: they assume a stable mapping from the underlying dynamics to the observed neural population activity throughout a task. As a result, they may fail to capture non-stationary processes in the brain that can lead to changing neuronal representations. A notable example of such a phenomenon is representational drift, which has been observed in various brain regions, ranging from the hippocampus (Ziv et al., 2013; Rubin et al., 2015) and posterior parietal cortex (Driscoll et al., 2017) to sensory regions such as V1 (Marks and Goard, 2021). Representational drift involves a gradual change in neural representations, where neuronal correlations with sensory and behavioral variables change across potentially different timescales, even when the observed behavior stays constant (Rule et al., 2019, 2020; Driscoll et al., 2022). Mounting evidence has also shown that this drift is not simply a result of experimental confounds such as recorded neuron turnover (Driscoll et al., 2022), suggesting the possibility that the activities of individual neurons change systematically while the underlying latent dynamics stay constant.

Traditionally, representational drift has been characterized as a long-term process, with studies focusing on changes that unfold across multiple sessions spanning days or weeks. Recent evidence, however, suggests that representational drift may occur on shorter timescales than previously reported, even within individual recording sessions lasting minutes to hours (Clarke et al., 2026). With a few notable exceptions (Carmena et al., 2005; Dahmen et al., 2022), shorter timescale changes in correlated patterns of neural activity have received limited attention in neuroscience, partly due to a lack of interpretable computational tools for detecting and quantifying subtle shifts in population-level structure at finer temporal resolutions.

Motivated by this, we introduce a new model class called the Stiefel Manifold Dynamical System (SMDS), which generalizes the LDS by allowing the mapping from latent states to observations to change over time. In the SMDS, emission matrices are constrained to lie on the Stiefel manifold—the space of all orthonormal matrices—and evolve smoothly along this manifold over trials. In contrast, the dynamics parameters are held fixed across trials. Consequently, SMDS models neural activity during a task as arising from a latent space with stable underlying dynamics, while allowing the mapping from this space to the observed activity to drift over trials. The orthonormal parameterization of the latents ensures this phenomenon is identifiable and allows us to quantify drift from changes in the emission matrices. To estimate the model parameters, we developed an variational expectation-maximization algorithm with extended Kalman smoothing.

We demonstrate the effectiveness of SMDS on both simulated and real neural datasets. In simulations, SMDS models outperformed LDS, which failed to recover the true dynamics and yielded lower held-out log-likelihood than SMDS in non-stationary settings. On neural recordings from macaques and rodents, SMDS again outperformed LDS on held-out data log-likelihood, crucially requiring fewer latent dimensions to capture the activity. Using the learned emission matrices from SMDS, we computed the Grassmann distance between them across trials and found a gradual drift in both datasets. SMDS also allowed us to decompose drift along individual subspace dimensions. Peak drifts ranged 13^◦^– 50^◦^ over 24 minutes in macaque primary motor cortex during a center-out reaching task (Clarke et al., 2026), and 37^◦^– 67^◦^ over 30 minutes in rodent anterior lateral motor cortex during a directional licking task (Chen et al., 2017). Notably, SMDS revealed less drift in dimensions encoding higher levels of neural and behavioral variance, suggesting that task-relevant information stays relatively more stable. Overall, we show that SMDS provides a powerful framework for tracking representational drift.^1^

## 2 Background

To develop our model for capturing representational drift, we require tools from both dynamical systems and differential geometry. We first introduce linear dynamical systems (LDS), a class of models for describing latent temporal structure in time series data. We then discuss identifiability issues in LDS and show how orthogonal parameterizations address them, motivating the use of Grassmann and Stiefel manifolds as the geometric framework for modeling how the mapping from latent dynamics to observations changes over time. Finally, we briefly review the extended Kalman filter (EKF) and extended Kalman smoother (EKS) in Appendix A, which enable approximate inference in nonlinear dynamical systems.

### 2.1 Linear Dynamical Systems

A standard linear dynamical system (LDS) describes observed data 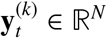 at timestep *t* of trial *k* as arising from a low-dimensional latent state 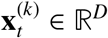. It consists of three main components: an initial state distribution, latent state dynamics, and an emission model.

The latent state, 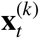, evolves over time according to a discrete-time linear Gaussian dynamics: 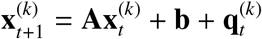, where **A** ∈ ℝ^*D*×*D*^ is the state dynamics matrix, **b** ∈ ℝ^*D*^ is the state bias vector, and 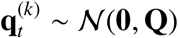 represents dynamics noise. We sample the initial state 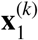 from an initial distribution 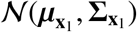. Finally, a linear mapping from the latent state generates the observations, 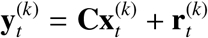, where **C** ∈ ℝ^*N*×*D*^ is the emission matrix, and 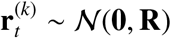 represents observation noise.

### 2.2 Identifiability in LDS

A key feature of LDS models is that there exists an equivalence class of parameters that all give rise to the same marginal distribution of observations. Specifically, for any invertible matrix **T** ∈ ℝ^*N*×*N*^, the transformed parameters 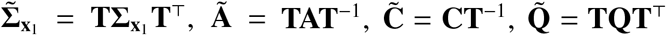 result in the same distribution of observations. Under this transformation, the latent states become 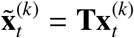.

We can reduce non-identifiability by constraining **C** to be orthonormal, **C**^⊤^**C** = **I**. Under this constraint, the only remaining transformations that preserve orthonormality are rotations within the latent space. Thus, this orthonormal parameterization identifies a unique subspace spanned by the emission matrix. This motivates the use of the Grassmann and Stiefel manifolds, which provide the natural geometric framework for working with such orthonormal matrices and their associated subspaces.

### 2.3 Grassmann and Stiefel Manifolds

The Grassmann manifold, denoted Grassmann(*N, D*), is the set of all *D*-dimensional subspaces of ℝ^*N*^ (Bendokat et al., 2024). Each point on the manifold is represented by a rank-*D* orthogonal projection matrix:

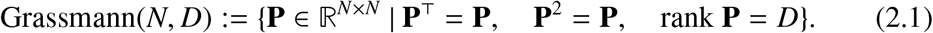

Each **P** ∈ Grassmann(*N, D*) corresponds to a unique *D*-dimensional subspace of ℝ^*N*^.

A given subspace can be represented by multiple different orthonormal bases—any rotation or reflection of the basis vectors spans the same subspace. The Stiefel manifold captures this collection of orthonormal basis (ONB) representations:

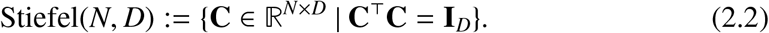

In our work, we parametrize the emission matrices **C**^(*k*)^ as elements of the Stiefel manifold, which allows us to track their evolution over trials while maintaining orthonormality. We then use the Grassmann distance to quantify drift by measuring changes in the underlying subspaces, which is invariant to rotations within those subspaces.

#### Displacements on the Stiefel manifold

To model how emission matrices evolve over trials while remaining orthonormal, we parametrize displacements on the Stiefel manifold using skew-symmetric matrices. At trial *k*, we construct a skew-symmetric matrix **B** ∈ ℝ^*N*×*N*^ as: 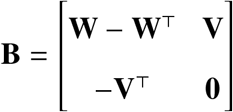 where **W** ∈ ℝ^*D*×*D*^ is a strictly upper-triangular matrix and **V** ∈ ℝ^*D*×(*N*−*D*)^ is arbitrary. The matrix **B** has two geometric roles: **V** governs motion orthogonal to the current subspace (inducing drift on the Grassmann manifold), while **W** governs rotations within the subspace (which do not change the underlying subspace).

We then map **B** to an orthogonal matrix using the Cayley transform, *f* (·):

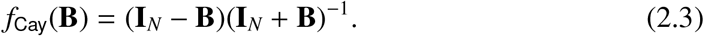

Note that we could alternatively map the skew-symmetric matrix **B** to an orthogonal matrix using the matrix exponential, but this is generally more costly and leads to a less well-behaved optimization problem (Wen and Yin, 2013).

#### Grassmann distance

The Grassmann distance quantifies how far two subspaces are from each other. Given ONB matrices **C**^(1)^, **C**^(2)^ ∈ Stiefel(*N, D*) and principal angles 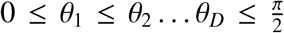 between their column spaces (Björck and Golub, 1973), the Grassmann distance is:

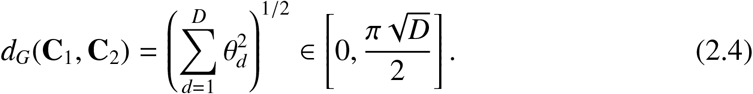

This corresponds to the geodesic distance—the shortest path between the two subspaces on the Grassmann manifold. We also define the normalized Grassmann distance as: 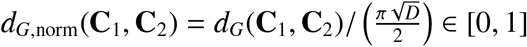.

## 3 Stiefel Manifold Dynamical System

Here we introduce the Stiefel Manifold Dynamical System (SMDS). We first define the generative model and then describe our inference procedure to infer latent trajectories of the orthonormal emission matrices on the Stiefel manifold. Finally, we describe a model selection strategy based on estimated marginal log-likelihood in Appendix C.

### 3.1 Model Definition

The Stiefel Manifold Dynamical System, as shown in Fig. 1, extends the LDS by allowing the emission matrices to vary across trials. We parameterize the evolution of the emission matrices through a latent “displacement” variable 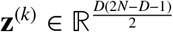, where *k* indexes the trial. We define the displacement as the concatenation of 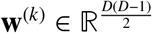 and the vectorization of **V**^(*k*)^ ∈ ℝ^*D*×(*N*−*D*)^:

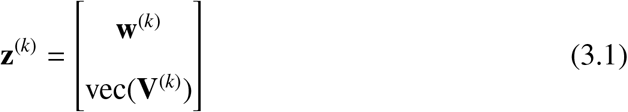

where **w**^(*k*)^, a vector consisting of the strictly upper-triangular entries of **W**^(*k*)^, governs rotations within the latent subspace, and **V**^(*k*)^ governs rotations that change the subspace itself. The displacement variables evolve across trials according to a random walk,

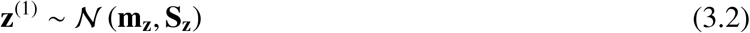

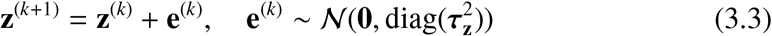

where 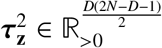 controls the rate of drift and is learned separately for each dimension so that each axis of the subspace can rotate at a different rate. The displacement variable defines a skew-symmetric matrix

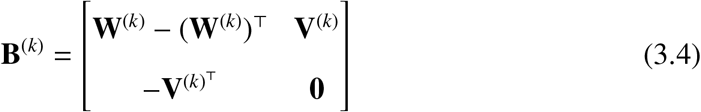

where **W**^(*k*)^ = triu(**w**^(*k*)^) ∈ ℝ^*D*×*D*^ is a strictly upper triangular matrix.

**Figure 1:**
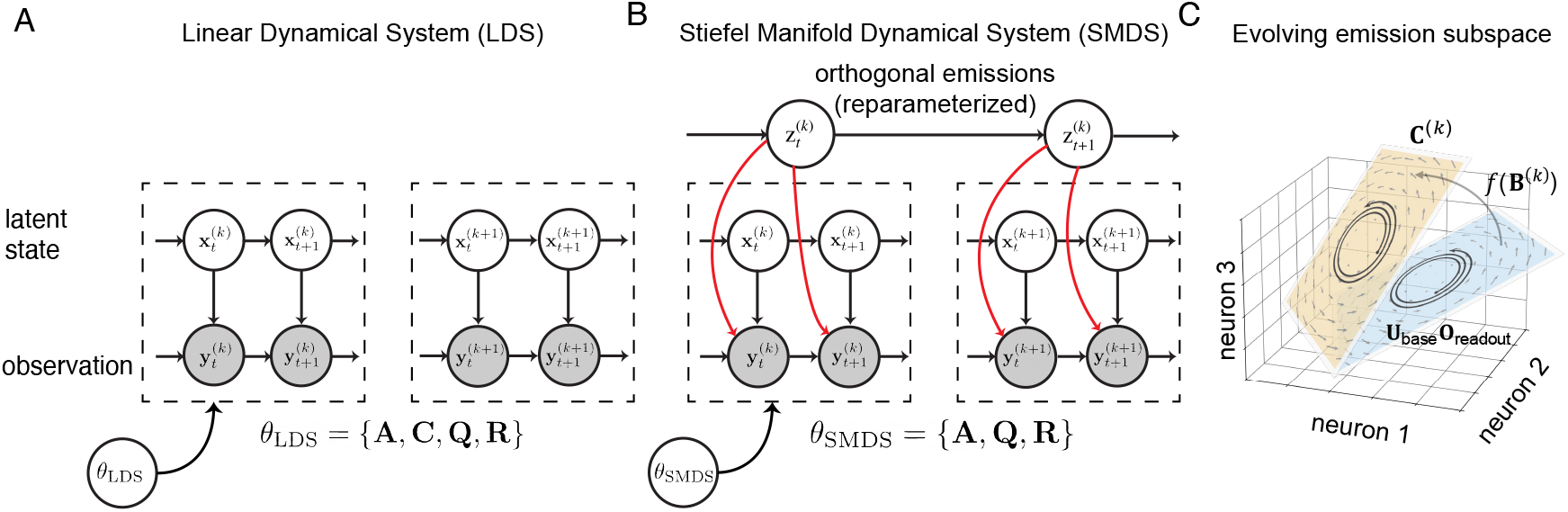
Stiefel Manifold Dynamical System. **(A-B)**: Graphical models of LDS and SMDS. At time *t* in trial 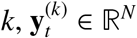 denotes the observed neural activity, and 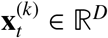 denotes the underlying low-dimensional latent state. SMDS, unlike a standard LDS, assumes that the emission subspace evolves smoothly over trials, parameterized by **z**^(*k*)^. **(C)**: Illustration of SMDS emission subspace evolution. **U**_base_**O**_readout_ is the fixed initial *D*−dimensional subspace in ℝ^*N*^. **z**^(*k*)^ captures the rotation at trial *k* relative to this, resulting in the rotated emission subspace **C**^(*k*)^ = **U**_base_ *f* (**B**^(*k*)^)**O**_readout_.

The emission matrix **C**^(*k*)^ for trial *k* is then defined as:

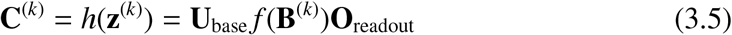

Where 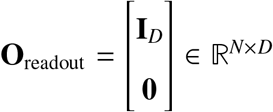, and **U**_base_ is a fixed set of coordinate axes spanning the ambient space ℝ^*N*^, in which the subspace lives and rotates. In practice, **U**_base_ is initialized using the principal components of the training data. We choose *f* (·) to be the Cayley transform, as described in Sec. 2.3. Thus, function 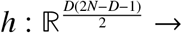 Stiefel(*N, D*) maps the displacement variable **z**^(*k*)^ to a point **C**^(*k*)^ ∈ Stiefel(*N, D*), with *f* (**B**^(*k*)^) defining the rotation matrix that transforms **U**_base_ at each trial *k*, and **O**_readout_ is a fixed matrix for reading out the first *D* columns (Fig. 1C).

The SMDS emission model is thus 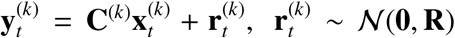, where the emission matrix varies over trials. The dynamics parameters {**A, Q**} are shared over trials, as in a standard LDS (Sec. 2.1). To complete the model specification, we place an Inverse-gamma prior on each entry of 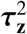, where the prior parameters are chosen via cross-validation (Appendix J).

### 3.2 Latent State and Parameter Inference

SMDS has two sets of latent variables: the within-trial latent states 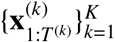 and the across-trial displacements 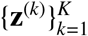, which parameterize the emission matrices. Computing the exact joint posterior over both sets of latent variables is intractable. Thus, we use approximate coordinate ascent variational inference under a structured mean-field factorization (Ghahramani and Hinton, 2000; Blei et al., 2017). We assume a factorized approximate posterior,

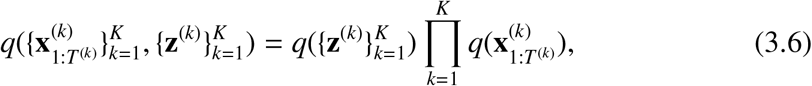

and alternate between updating each factor while holding the other fixed, interleaved with M-step updates of the model parameters 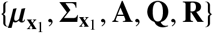.

#### Updating 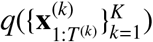: Kalman smoothing

Given the current estimate of the displacements, we fix the emission matrices at 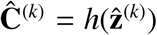, where 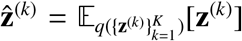 is the posterior mean. While it is suboptimal to condition on only a point estimate of the emission matrix rather than the propagating uncertainty from the variational factor, 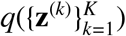, this approach is much more straightforward. Given a point estimate, the update for the within-trial latent states reduces to a standard LDS, and we update the factor 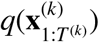 using exact Kalman smoothing for each trial *k*.

#### Updating 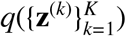: Extended Kalman smoothing

Given the current 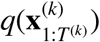 for *k* = 1, …, *K* from the previous step, we update the approximate posterior over the displacements. With the structured mean field approximation, the optimal update is,

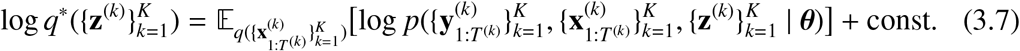

As derived in Appendix B, the expected log joint probability on the right-hand side is equivalent to a state space model with linear Gaussian dynamics (from the prior on 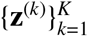) and *nonlinear* Gaussian emissions,

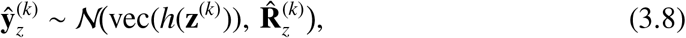

where 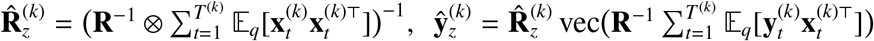, and, ⊗ denotes the Kronecker product. Again, the exact coordinate ascent update is challenging to compute. Instead, noting that the emission function vec(*h*(·)) is nonlinear, we propose to use an extended Kalman smoother (EKS) to compute an approximate Gaussian posterior 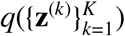, analogous to previous work (Zoltowski et al., 2020).

#### M-step

Given the approximate posteriors 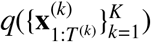 and 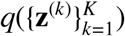, we update the model parameters. The dynamics and observation noise parameters 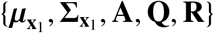 are updated in closed form, as in a standard LDS. The displacement prior parameters {**m**_**z**_, **S**_**z**_} and drift rates 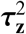 are updated using the posterior statistics from 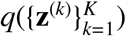. We find that the quality of the first-order approximation used in the EKS depends on the magnitude of 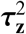 (see details in Appendix D). As such, we clip 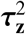 after each M step, where the clipping threshold is also a tunable hyperparameter (Appendix J).

## 4 Results

We first compare SMDS to LDS on synthetically generated non-stationary data. Next, we apply SMDS, LDS, and conditionally linear dynamical systems (CLDS) (Geadah et al., 2025) to analyze real macaque neural recordings during a center-out reaching task and rodent two-photon calcium imaging data during a whisker-based directional licking task. CLDS uses Gaussian process priors to allow model parameters to vary smoothly with observed covariates such as reaching directions. We adapt CLDS to model representational drift by setting the covariate to the trial block index and allowing only the emission matrix to vary across trials, while keeping the dynamics parameters fixed (see Appendix F for details and Appendix J for hyperparameters).

### 4.1 Application to simulated data

We begin with a toy example to visually demonstrate the advantages of SMDS over a standard LDS. We simulated data from SMDS with latent dimension *D* = 2 and observation dimension *N* = 10. The simulated dataset consisted of a total of 750 trials, with 30 timesteps per trial, where the emission matrices smoothly rotated over trials (see details in Appendix G.1). We fitted both LDS and SMDS across a range of state dimensions, *D* ∈ [1, 10], and evaluated model performance based on held-out data log-likelihood (Fig. 2A).

**Figure 2:**
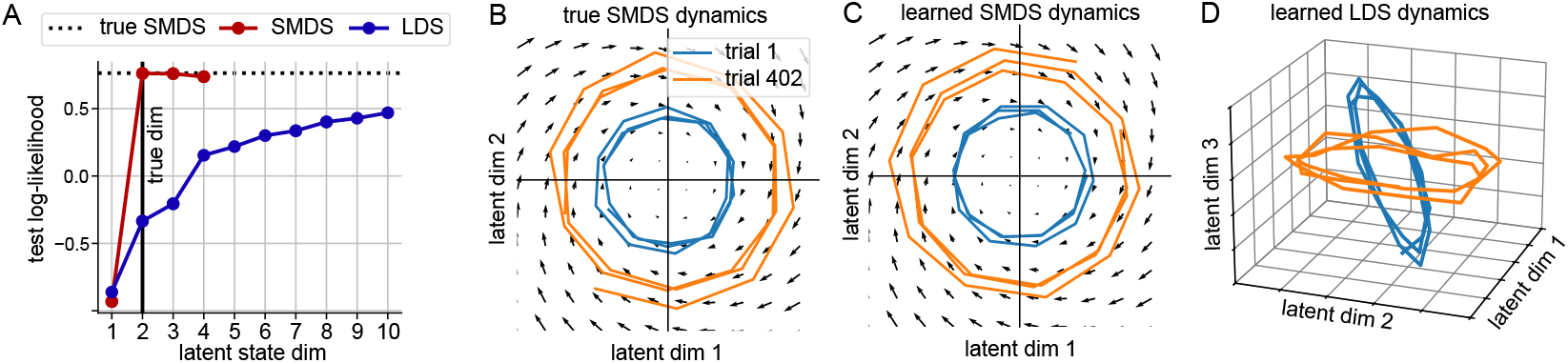
Simulated data experiment 1. **(A)**: Held-out log-likelihood on data generated from an SMDS with *D* = 2, *N* = 10. SMDS outperforms standard LDS and recovers the true state dimension. **(B–D)**: SMDS fitted to this data with *D* = 2 recovers ground truth dynamics, while LDS requires higher-dimensional states.

We found that the test log-likelihood of SMDS saturated at the ground truth latent dimensionality, and that SMDS achieved higher test log-likelihood than LDS. The test log-likelihood of LDS continued to increase beyond the true latent dimensionality, failing to identify the correct dimensionality. We also visualized the true and learned smoothed latent trajectories for two example trials (Fig. 2B). While SMDS recovered the true two-dimensional dynamics up to a rotation (Fig. 2C), the LDS with *D* = 3 learned trial-specific shifted dynamics (Fig. 2D) to compensate for drift in the observation space. Although the test log-likelihood for LDS kept increasing, we chose *D* = 3 for visualization purposes. We show in Appendix G.2 that SMDS also recovers ground-truth dynamics and drift in higher-dimensional settings (*D* = 8, *N* = 24), including per-dimension drift rates.

### 4.2 Modeling macaque neural data during center-out reaching task

We applied SMDS to neural recordings from Clarke et al. (2026),^2^ where a macaque performed a center-out reaching task (Fig. 3A) The dataset comprised 96-channel Utah array multiunit activity from M1 spanning 750 trials from a post-lesion session (Fig. 3B). Trials were grouped in blocks of eight, each containing all conditions in random order. We binned neural activity in 25 ms bins and performed our analysis on activity from -50 ms to 450 ms relative to movement onset.

**Figure 3:**
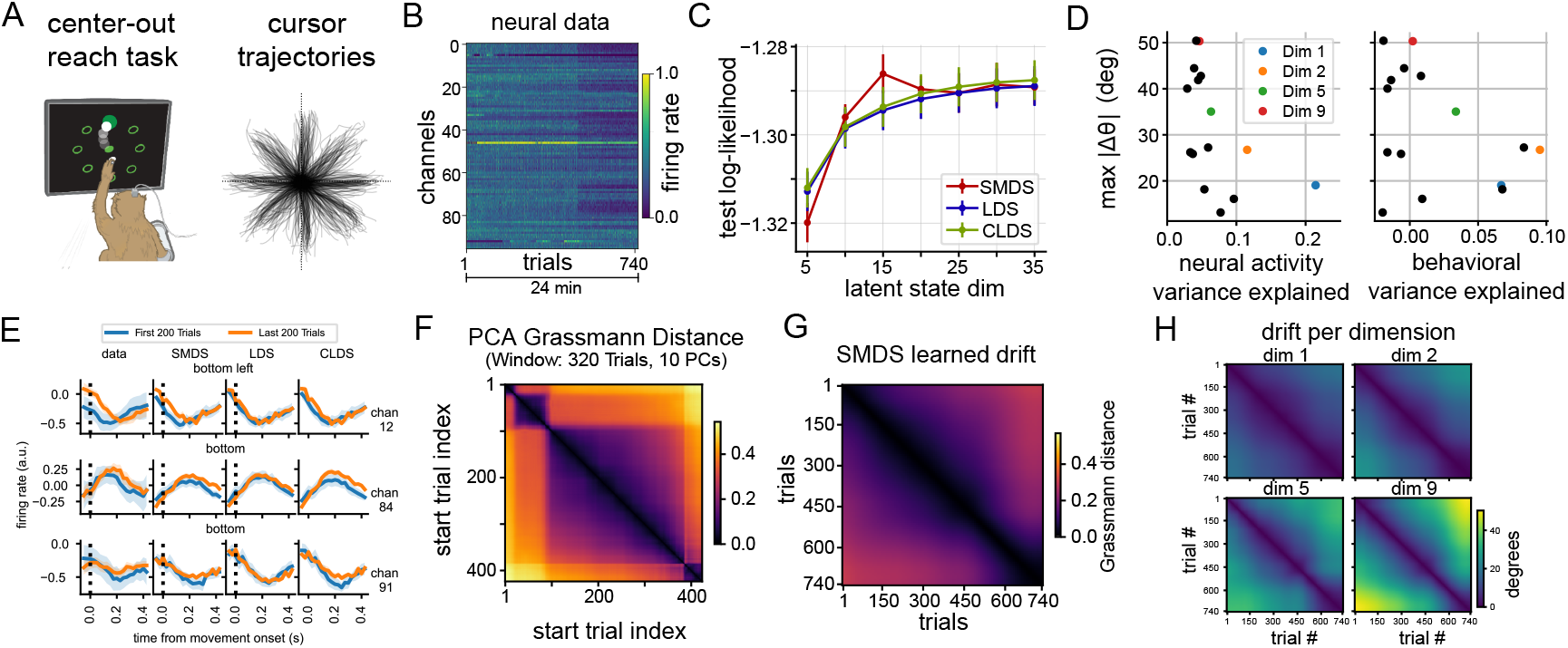
SMDS drift analysis on neural recordings from a macaque performing a center-out reaching task. **(A)** Task schematic and example center-out reach trajectories. **(B)** Raster plot of neural activity from a single session. The dataset includes recordings from 96 channels, with a total of 744 trials. **(C)** Held-out log-likelihood comparison. SMDS outperforms LDS and CLDS using fewer latent dimensions. **(D)** Peak drift magnitude during the session vs. explained variance per dimension: neural (left) and behavioral (right). Axes that account for more neural variance drift less, whereas less informative axes drift more **(E)** PSTHs from early and late trials for a subset of channels under the “bottom” and “bottom left” reach conditions, aligned to movement onset, showing evidence of representational drift (left to right: data, SMDS, LDS, and CLDS predictions). **(F)** Normalized Grassmann distance matrix computed using the top 10 principal components over a sliding window of 320 trials. **(G)** Overall drift inferred by SMDS, computed using Grassmann distance between pairs of learned emission matrices. **(H)** Estimated drift per dimension, computed in degrees.

We found evidence of representational drift through both visual investigation and principal component analysis of the raw data. Fig. 3B & E (left) show average firing rates over trials and peri-stimulus time histograms (PSTHs), respectively, for three example channels. Both firing rates and PSTHs change between early and late trials. Additionally, the normalized Grassmann distance between subspaces spanned by the top 10 principal components (PCs), computed using sliding windows of 320 trials, revealed subspace rotation indicative of drift (Fig. 3F).

We fitted SMDS, LDS, and CLDS using 5-45 latent dimensions across five train-test splits, holding out 12 random blocks per split. We allowed the emission matrices to evolve across blocks in SMDS and CLDS (see Appendix H for all hyperparameter selection details). SMDS outperformed both LDS and CLDS in terms of marginal log-likelihood on held-out blocks, saturating at a smaller latent dimensionality (Fig. 3C), highlighting its ability to model non-stationary data effectively. Predicted PSTHs from SMDS also closely matched real data, as shown in Fig. 3E.

To characterize the drift learned by SMDS, we computed the normalized Grassmann distance between emission subspaces across blocks, revealing a gradual drift throughout the session (Fig. 3G). Decomposing the drift along individual latent dimensions, we found that dimensions explaining more variance in the neural data (Fig. 3D left) and those that are behaviorally significant—in terms of predicting cursor velocity—exhibited less drift (Fig. 3D right). Conversely, less informative dimensions showed more drift. Visualization of pairwise drift between blocks for selected latent dimensions (Fig. 3H) revealed varying patterns. Some dimensions (e.g., 9) rotated up to 50^◦^ and showed structured drift patterns similar to the PC subspace drift (Fig. 3D & F), while others (e.g., 1) exhibited more modest drift around 19^◦^.

Thus, these findings reveal gradual representational drift in macaque M1 during the reach task. However, this drift is not uniform across all latent dimensions. Instead, it appears to preferentially affect less neurally and behaviorally significant dimensions, potentially to maintain stable representations of task-relevant information over time.

### 4.3 Modeling rodent neural data during a directional licking task

We next applied SMDS to a two-photon calcium imaging dataset from Chen et al. (2017), where a rodent performed a whisker-based object localization task with a delayed, directional licking response (Fig. 4A). The dataset included 126 correct trials from a total of 132 regions of interest (ROIs) in the anterior lateral cortex (ALM) and medial motor cortex (MM), imaged at 14Hz. We analyzed 7 seconds of data around the go cue per trial (98 time bins/trial). As with the macaque dataset, we saw evidence of changes in neural representations within the session through two analyses of the raw data: (1) visual inspection of trial-averaged responses (Fig. 4B) and PSTHs during early and late trials (Fig. 4E left), and (2) pairwise normalized Grassmann distance between the subspaces spanned by the top 7 PCs, using sliding windows of 32 trials (Fig. 4F).

**Figure 4:**
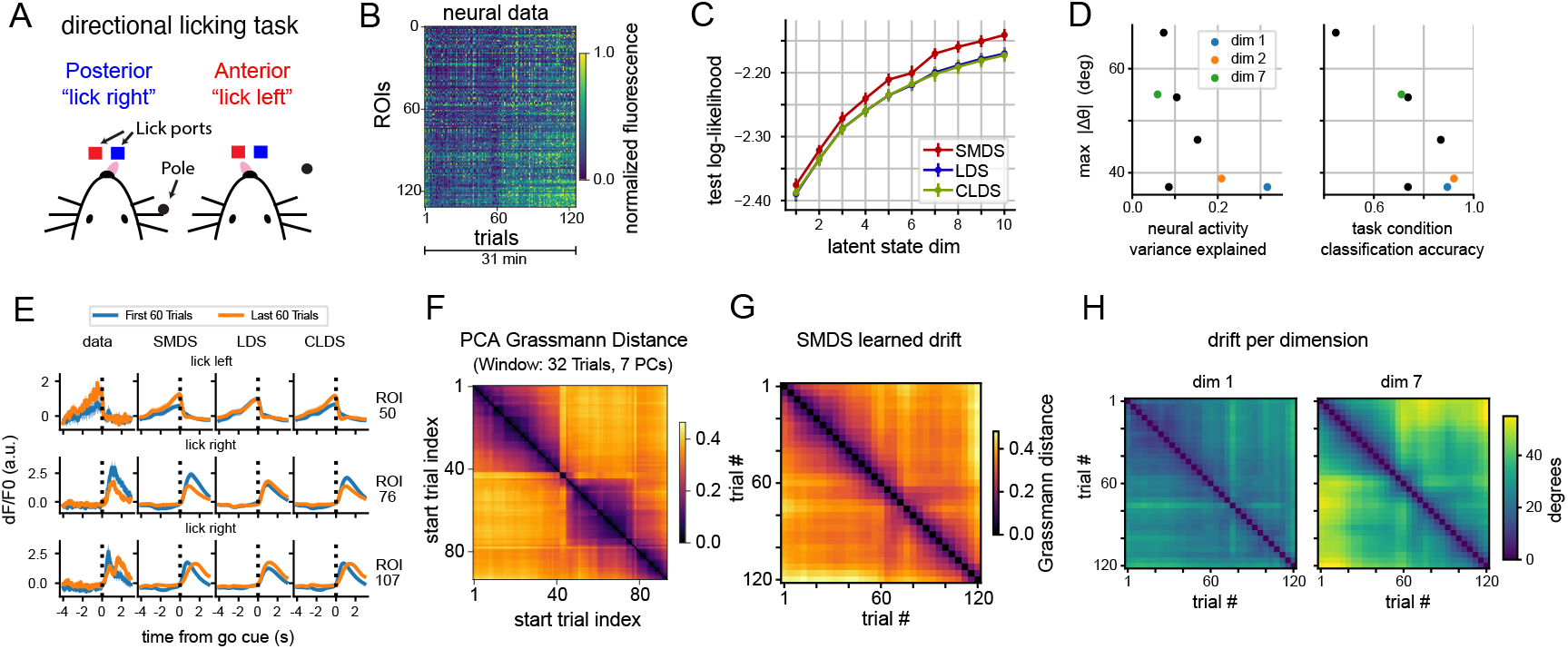
SMDS drift analysis on a rodent neural dataset during a directional licking task. **(A)** Task schematic. The rodent licks either the left or right port based on the pole’s location. **(B)** Two-photon calcium imaging fluorescence from a single session. The session includes recordings from 132 ROIs over approximately 31 minutes, consisting of 155 trials. We analyzed 126 correct trials with no early licks. **(C)** Held-out log-likelihood comparison showing that SMDS consistently outperforms standard LDS. **(D)** Peak drift magnitude vs explained neural variance per dimension (left), and trial condition classification accuracy (right). Axes that account for more neural and behavioral variance drift less. **(E)** PSTHs from early and late trials for a subset of ROIs, aligned to the “go” cue (left to right: data, SMDS, LDS, and CLDS predictions) **(F)** Normalized Grassmann distance matrix computed using the top 7 principal components over a sliding window of 32 trials. **(G)** Overall drift inferred by SMDS, computed using Grassmann distance between pairs of learned emission matrices. **(H)** Estimated drift per dimension, computed in degrees.

We then fitted SMDS, LDS, and CLDS using 1-10 latent dimensions across three train-test splits (see Appendix I for hyperparameter details). For SMDS and CLDS, we grouped trials into blocks of four, allowing emissions to evolve over blocks. Once again, SMDS outperformed both LDS and CLDS in terms of marginal log-likelihood on held-out blocks and required fewer latent dimensions (Fig. 4C). We also visualized PSTHs predicted by SMDS (Fig. 4E) and found that they closely match the data.

We then analyzed the learned subspace changes along individual latent dimensions. Similar to the macaque dataset, dimensions that explained more neural variance and better classified trial conditions exhibited smaller peak drift (Fig. 4D), reinforcing our hypothesis that representations of task-relevant information tend to remain more stable. Visualization of pairwise normalized Grassmann distances between the learned emission subspaces over blocks revealed a gradual drift over the session (Fig. 4G). When visualizing drift for selected latent dimensions, some dimensions exhibited high and structured drift—for example, dimension 7 rotated by over 55^◦^ and showed structured drift patterns similar to the PC subspace drift (Fig. 4F)—while others, such as dimensions 1 and 2, showed slower drift with a peak around 37^◦^ (Fig. 4H). These findings emphasize that different latent dimensions drift at varying rates and patterns, where neurally and behaviorally significant dimensions tend to be more stable.

## 5 Related Work

### Subspace Tracking

Subspace tracking, the problem of dynamically estimating the subspace of streaming data, has a long history, from Oja’s algorithm (Oja, 1982) to modern methods handling high-dimensionality, limited memory, and data corruption (Yang, 1995; Balzano et al., 2018; Narayanamurthy and Vaswani, 2018a,b; Bienstock et al., 2022). Recent approaches model subspace evolution on manifolds: Sasfi et al. (2024) used Grassmannian gradient descent for subspace tracking in LDS, Blocker et al. (2023) estimated piecewise geodesic subspace trajectories, and Srivastava and Klassen (2004) and Rentmeesters et al. (2010) used particle filtering on the Grassmann manifold. Related Stiefel manifold methods employ matrix Langevin noise models with particle filtering or iterative Bayesian updates (Tompkins and Wolfe, 2007; Bordin and Bruno, 2020; Chikuse, 2006). SMDS is related to these approaches in that it allows the emission matrices to evolve on a Stiefel manifold, but differs in that it performs inference using an extended Kalman smoother.

### Analysis of Nonstationary Neural Data

Prior work addresses neural nonstationarity through alignment methods that reveal stable structure across sessions (Gallego et al., 2020), switching models that capture discrete transitions between dynamical regimes (Ghahramani and Hinton, 2000; Linderman et al., 2017; Glaser et al., 2020; Costacurta et al., 2022; Lee et al., 2023), sparse decompositions of nonstationary dynamics (Mudrik et al., 2024), and condition-dependent subspace models where parameters vary smoothly with task variables (Nejatbakhsh et al., 2023; Geadah et al., 2025). Our focus is to model changes in the observed neural representation space, while the underlying dynamics stay constant.

## 6 Discussion

We introduced the Stiefel Manifold Dynamical System (SMDS), a novel class of probabilistic state space models that learns neural dynamics while accounting for representational drift. SMDS constrains its emission matrices to be orthonormal and allows them to evolve smoothly over trials on the Stiefel manifold. On simulated data, SMDS recovered true latent dynamics and outperformed LDS, which failed to account for non-stationarity. On macaque and rodent neural recordings, SMDS revealed gradual drift occurring over a single experimental session. Our approach also revealed that dimensions encoding higher neural and behavioral variance drifted less relative to others, thus maintaining stable task-relevant representations. These findings underscore SMDS’s potential for understanding neural dynamics and their evolution over time and open new avenues for investigating sources and causes of representational drift.

Several directions remain for future work. While SMDS assumes smooth drift over trials, it may struggle to capture abrupt or event-driven drift in neural activity. Future work could exploit discrete switching transitions in drift rates to address this. Next, SMDS could be extended to relax the shared dynamics assumption, thus allowing the model to learn changes in the underlying neural computations over time, along with drift in the observation space. Finally, SMDS models the evolution of neural subspaces independently of behavior. Jointly modeling neural and behavioral data may enable SMDS to learn a time-varying mapping between the two. Despite these limitations, SMDS provides a principled framework for quantifying representational drift and understanding neural dynamics in non-stationary settings.

## Acknowledgments and Disclosure of Funding

HDL is supported by the Ketterer-Vorwald Stanford Interdisciplinary Graduate Fellowship. AJ is supported by the Wu Tsai Interdisciplinary Postdoctoral Fellowship. We also thank the members of the Linderman Lab and Brain Interfacing Lab for their support and feedback throughout the project. MS is supported by NSF GRFP DGE-1656518 and an NIH Diversity Supplement R01NS130789-S1 (to PN for MS). SEC is supported by SoM Dean’s Postdoctoral Fellowship and a Stanford HAI Seed Research Grant (SEC & PN). PN is supported by grants from the NIH (R01NS123517 R01NS130789 U19NS118284) and the Wu Tsai Neurosciences Institute. SWL is supported by grants from the NIH BRAIN Initiative (U19NS113201, R01NS131987, R01NS113119, & RF1MH133778), the NSF/NIH CRCNS Program (R01NS130789), and the Sloan, Simons, and McKnight Foundations. We thank the members of the Linderman Lab and the Brain Interfacing Lab for feedback throughout this project. We thank Kimberly Chin for administrative support.

## Appendix

### A Extended Kalman Filters and Smoothers

While Kalman filters and smoothers are well-suited for linear systems, they are not directly applicable when either the dynamics or the emission models are nonlinear. Here we briefly describe the extended Kalman filter (EKF) and extended Kalman smoother (EKS), which are recursive algorithms for approximate inference in systems with nonlinear dynamics or nonlinear observation models (Särkkä, 2013).

The EKF linearizes the dynamics and observation functions around the current state estimate at each timestep using their first-order Taylor approximation, thus allowing a local application of the standard Kalman filter update. Specifically, given a nonlinear state transition function 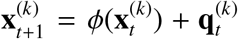 and emission model 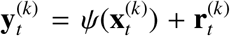, the EKF computes the Jacobians of *ϕ* and *ψ* at the current estimate and propagates the mean and covariance of the state forward in time. These linearizations yield an approximate Gaussian posterior over the latent state at each timestep, enabling efficient filtering in otherwise intractable nonlinear models. Analogous to Kalman smoothing, the EKS builds on the EKF by incorporating future observations to refine the latent state estimates.

### B Derivation of the Displacement Update in SMDS

We derive the update for the approximate posterior over the displacements 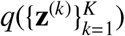 used in Sec. 3.2. As described in Sec. 3.2, we assume a mean-field factorization

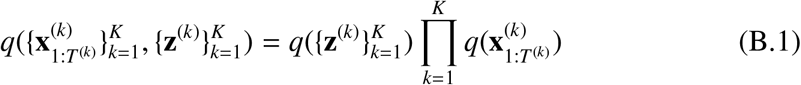

and optimize the ELBO by approximate coordinate ascent, alternating between the two factors. The ELBO is

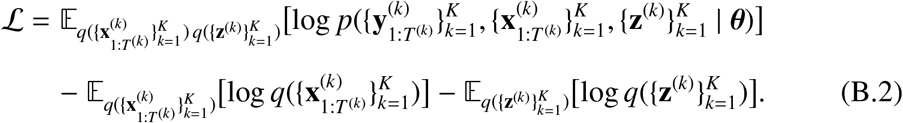

Holding 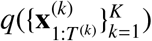 fixed and optimizing over 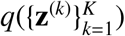, the terms that depend on 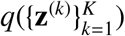 are

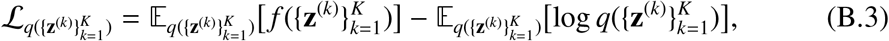

rwhere 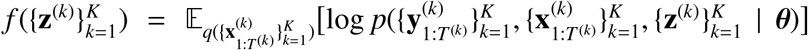. Defining 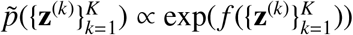, we can rewrite (B.3) as

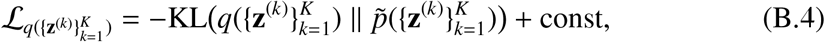

which is maximized when 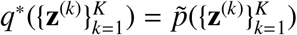

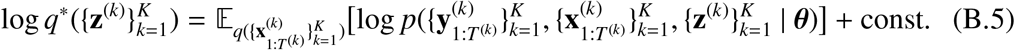

Expanding the joint and dropping terms that do not depend on 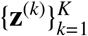,

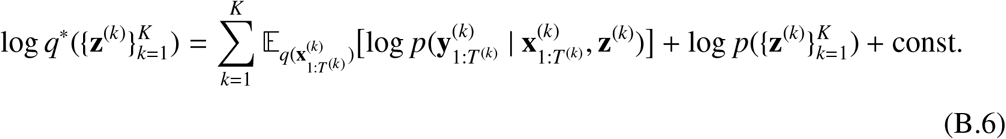

Below we show that (B.6) takes the form of a state-space model with a nonlinear emission function, allowing us to approximate *q*^∗^ with an extended Kalman smoother. We begin by simplifying the first term.

#### Per-trial emission log-likelihood

We begin with the emission log-likelihood for a single trial *k*, conditioned on the latent states and the displacement:

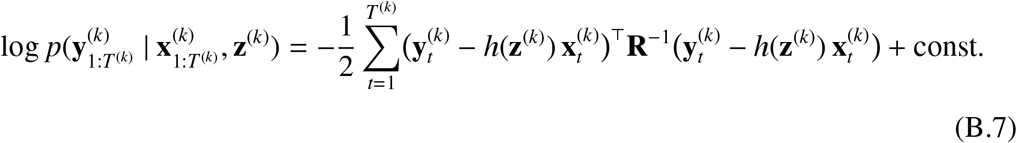

Expanding the quadratic form and dropping terms that do not depend on **z**^(*k*)^,

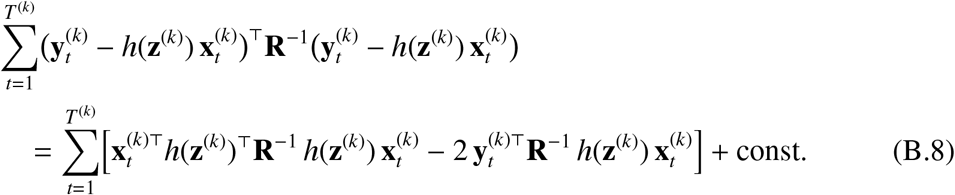

#### Vectorized form

Using the identities tr(**XAX**^⊤^**B**) = vec(**X**)^⊤^(**B** ⊗ **A**) vec(**X**) (for symmetric **A** and **B**) and tr(**A**^⊤^**B**) = vec(**A**)^⊤^vec(**B**), we rewrite (B.8) in terms of vec(*h*(**z**^(*k*)^)):

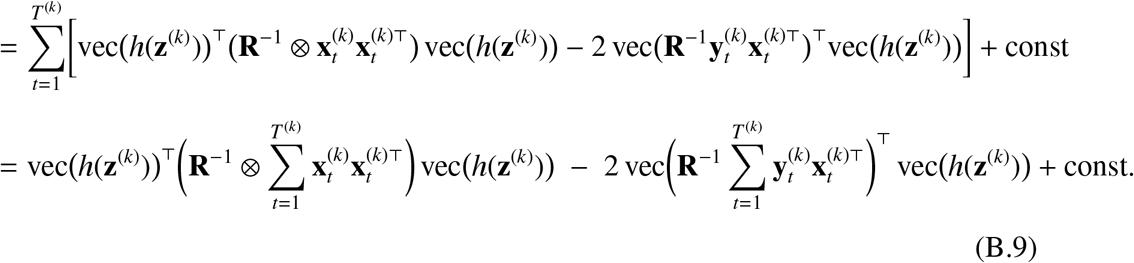

#### Completing the square

Taking expectations under 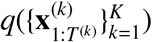 and completing the square in vec(*h*(**z**^(*k*)^)), the expected emission log-likelihood (B.7) is equivalent (up to a constant) to

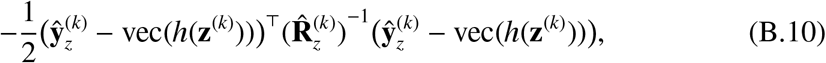

where

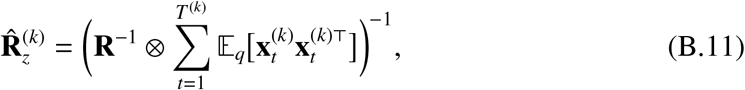

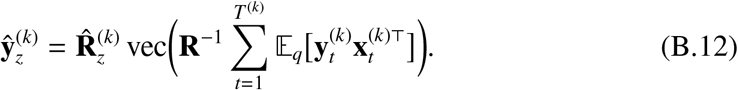

Here, ⊗ denotes the Kronecker product.

#### State-space model for the displacements

Substituting (B.10) into (B.6), we can interpret the right-hand side as the log joint of a state-space model with linear dynamics (from the random walk prior log 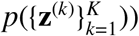 and a nonlinear emission function:

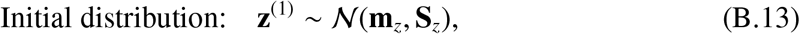

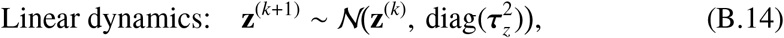

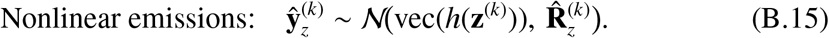

Concretely, since 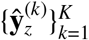 are fixed quantities (computed from the data and the current 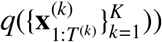, we have

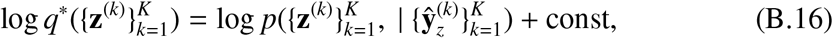

We approximate *q*^∗^ using an extended Kalman smoother, yielding an approximate Gaussian posterior 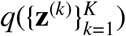. The posterior mean 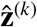 gives the estimated emission matrix 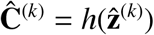.

### C Computing Marginal Log-likelihood of Data under an SMDS

To evaluate and compare the performance of SMDS with a standard LDS, we approximate the marginal log-likelihood of held-out data given the training data to test the model’s ability to generalize to unseen data. Approximating the marginal log-likelihood for an SMDS model requires marginalizing over both latent states and displacements. Naïvely, this would require memory of 𝒪(*N*^2^*T* ^2^) where *N* is the data dimensionality and *T* is the length of a trial (Appendix C.1). Thus, for computational and memory efficiency, we formulated an equivalent representation of SMDS in an augmented latent space that jointly includes the latent states and displacements (Appendix C.2). This eliminates the need to store statistics for each time point in memory and allows us to use an EKF in the augmented space and obtain marginal log-likelihood estimates efficiently.

#### C.1 Block-wise Approach

Let 𝓎 ∈ ℝ^*K*×*T*×*N*^ be the tensor denoting the data. Here, we describe the naïve approach to approximate the marginal log-likelihood. Omitting the dependence on the learned parameters for simplicity, we can approximate it as follows:

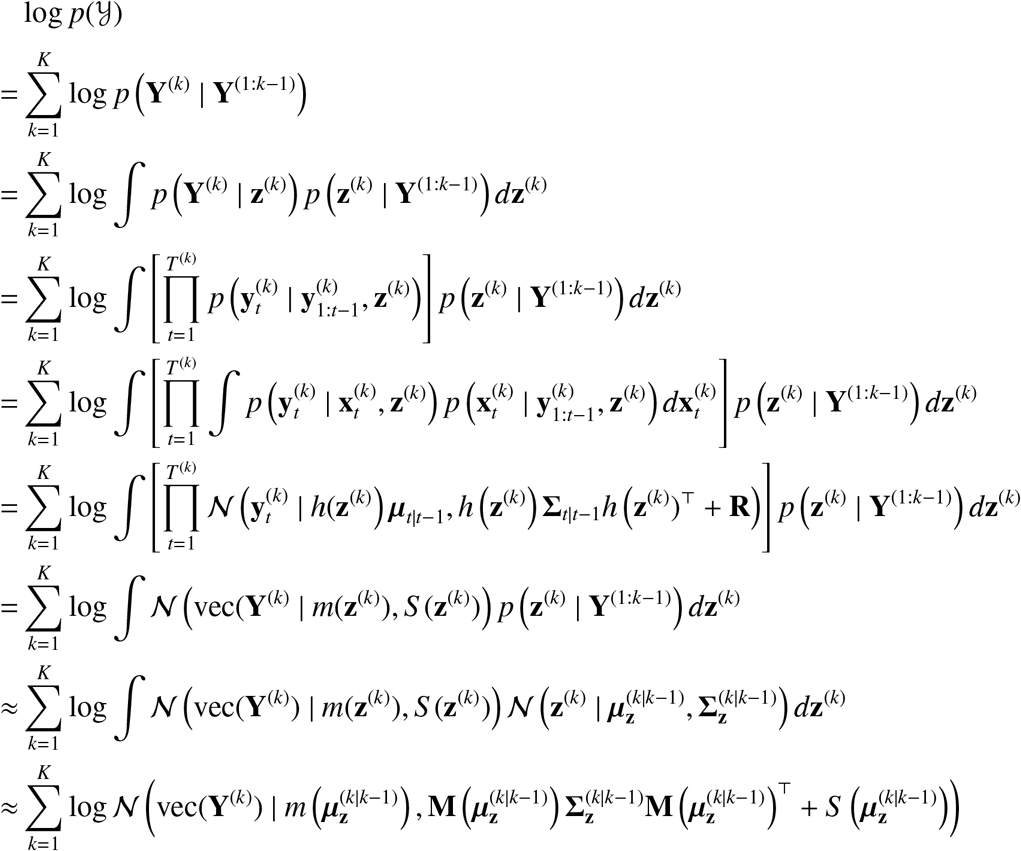

where

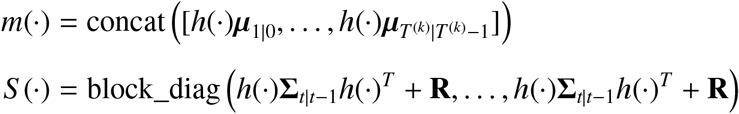

and **M**(·) is the Jacobian of *m*(·). This block-wise approach requires materialization of the matrix *S* (·) ∈ ℝ^*NT*×*NT*^, which quickly becomes intractable for large *T* (i.e., long trials or blocks of trials).

#### C.2 Augmented Space Approach

We denote the augmented state for trial *k* and timestep *t* as

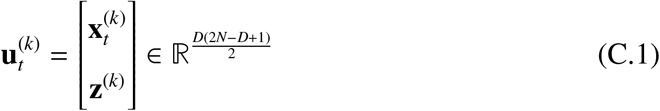

where 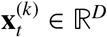 and 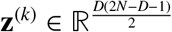 . In this augmented space, the within-trial dynamics is:

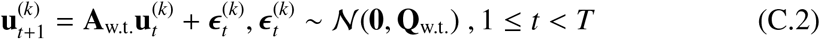

Where 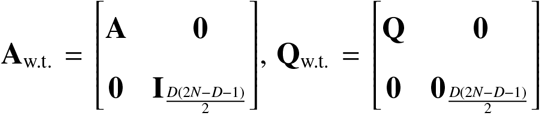, and *T* is the length of a trial. At trial boundaries, we reset the latent states 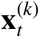 to their initial distribution, while the displacement **z**^(*k*)^ evolves per their random walk dynamics, resulting in the following across-trial dynamics:

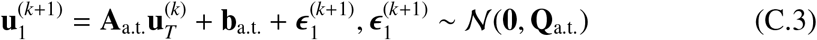

where 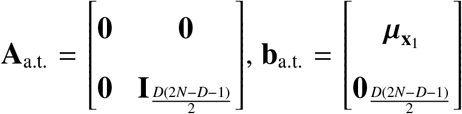, and 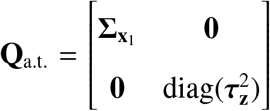. Finally, we can express the mapping from this space to the observations as:

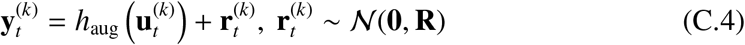

*h*_aug_ is defined as a nonlinear transformation:

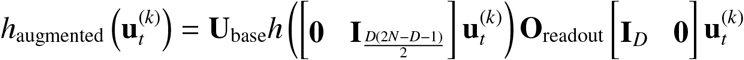

where 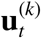 is the augmented state at time *t* of trial *k*. **U**_base_ and **O**_readout_ are defined in Sec. 3.1. Intuitively, this function first splits the augmented state 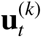 into 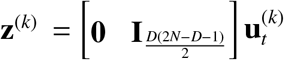 and 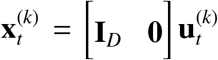 and then applies the function *h*(·) defined in Sec. 3.1. Using this formulation, we use an EKF in the augmented state space to obtain marginal log-likelihood estimates for SMDS.

### D First-order Approximation of the SMDS Emission Model

Here we explore how the extended Kalman filter and extended Kalman smoother’s first-order approximation of the nonlinear emission model of SMDS changes with the drift parameter τ^2^. Consider *N* = 2 and *D* = 1. Let **z** ∼ 𝒩(**0**, diag(τ^2^)) and **y** ∼ 𝒩(*h*(**z**), *σ*^2^**I**_*N*_), where *h*(·) is the nonlinear function that maps a displacement to an emission matrix, as defined in Sec. 3.1. Note that **z** ∈ ℝ^1×1^ and 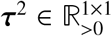. In addition, let **U**_base_ = **I**_*N*_. Then, we have *h*(**z**) = *f*_Cay_ (**B**)**O**_readout_, where 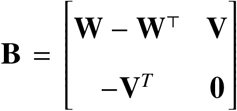 and 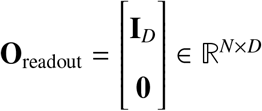. Note that for *D* = 1, **W** is a 1 × 1 zero matrix. For various values of τ^2^, we sample 1,000 **V**’s to get samples of **y**’s. We compare the sample mean and covariance against their first-order approximations, which can be computed by *h*(**0**) for the mean and *H*(**0**)diag(τ^2^)*H*(**0**)^*T*^ + *σ*^2^**I**_*N*_ for the covariance, where *H*(·) is the first-order Taylor approximation of the function *h*(·). We set *σ*^2^ = 0.01 for the visualizations in Fig. 5. As we can see, the first-order Taylor approximation error increases as τ^2^ increases. Motivated by this observation, we clip τ^2^ after each M step in SMDS, such that it stays below a the clipping threshold, which is a tunable hyperparameter.

**Figure 5:**
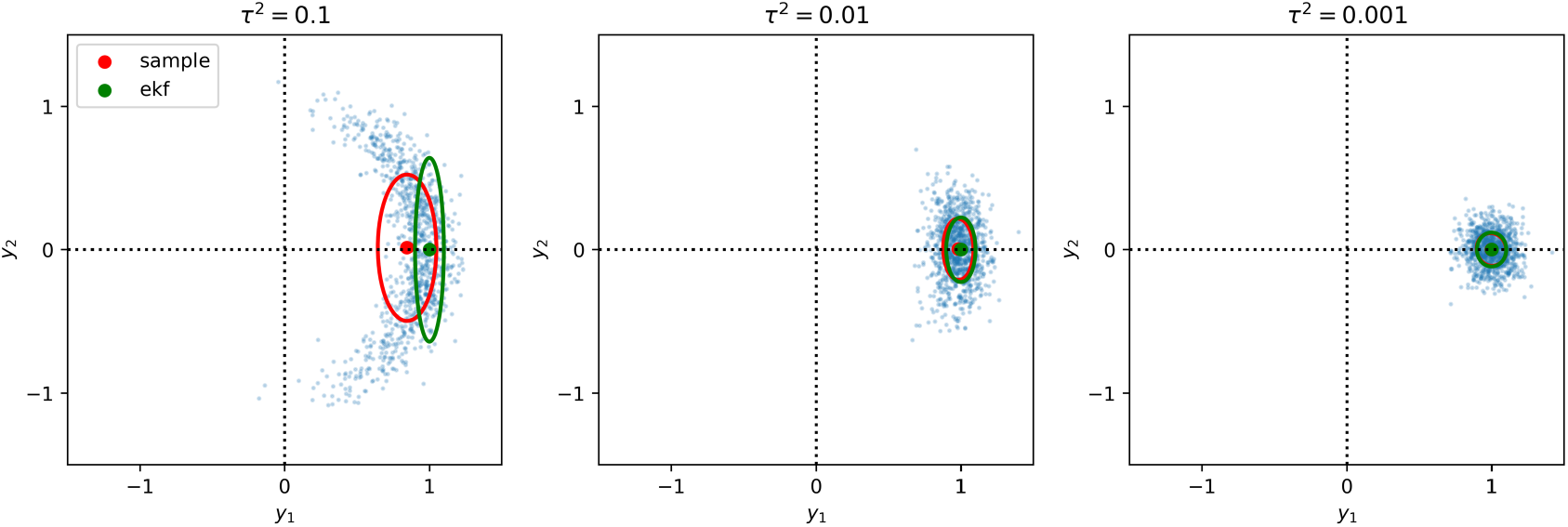
First-order Taylor approximation error of the nonlinear emission model of SMDS increases as τ^2^ increases. Sample mean and covariance versus first-order Taylor approximation of the mean and covariance of the emission model of SMDS, for various levels of τ^2^.

### E Max |Δ*θ*| (Peak Drift) Computation

Here we describe how we compute the maximum |Δ*θ*|. First, for each dimension *d*, we compute the cosine similarity for each pair of trials:

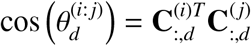

for *i, j* ∈ {1, …, *K*}. Here, 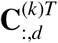 denotes the *d*-th column of the emission matrix for trial *k*. Note that we can omit the normalization since 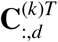 is a unit vector. We then take the *arccos* of the cosine similarity to get 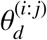, the amount of rotation of the *d*-th dimension in radians from trial *i* to trial *j*. After converting 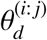 into degrees, we compute the maximum of 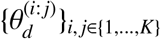 to get the maximum |Δ*θ*| across all trials. Intuitively, this metric measures the maximum separation that ever occurred for dimension *d* throughout the session.

### F Conditionally Linear Dynamical Systems

In the original Conditionally Linear Dynamical System (CLDS) formulation (Geadah et al., 2025), the dynamics and emission parameters vary smoothly as functions of an observed covariate **u** (e.g., heading direction or reach angle) through Gaussian process (GP) priors. To adapt CLDS for modeling representational drift, we set the covariate to the normalized trial block index, 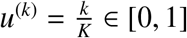, where *K* is the total number of blocks, and restrict non-stationarity to the emission matrix alone. Concretely, the dynamics parameters {**A, b, Q**} and the observation noise **R** are shared across all trials, while the emission matrix varies over blocks:

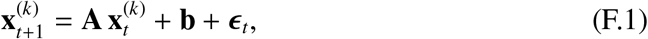

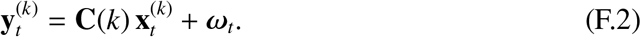

Each entry of **C**(*k*) is parameterized via a truncated Fourier feature expansion approximating a squared-exponential GP prior:

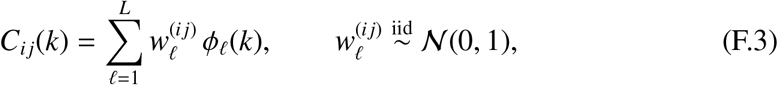

where 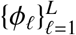 are Fourier basis functions on a torus of period *p*, designed to approximate a squared-exponential GP with lengthscale *κ* and scale *σ*. Following the recommendation in the CLDS codebase^3^, the period is set to *p* = 1 + 6*κ*, placing the periodic boundary well outside the covariate domain [0, 1] and rendering the prior effectively non-periodic over the observed trials. The lengthscale *κ* governs how rapidly the emission matrix evolves across trials, playing a role analogous to the inverse of 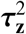 in SMDS.

Note that, unlike SMDS, CLDS does not constrain the emission matrix to be orthonormal and thus does not enforce identifiability of the latent subspace. The GP hyperparameters {*κ, σ, L*} were selected via grid search over held-out log-likelihood. See Table 2 in Appendix J for the values searched.

**Table 1:** Outline of the key hyperparameters in LDS and SMDS.

| Hyperparameter | Description | Applicable Models | Permissible Values |
| --- | --- | --- | --- |
| $D$ | Dimension of the latent state space. | LDS, SMDS, CLDS | $\mathbb{Z}_{\geq 1}$ |
| $\alpha$ | Shape of the Inverse-gamma prior on $\tau^2$. | SMDS | $\mathbb{R}_{>0}$ |
| $\beta$ | Scale of the Inverse-gamma prior on $\tau^2$. | SMDS | $\mathbb{R}_{>0}$ |
| Initial $\tau^2$ | Initialization value of $\tau^2$. | SMDS | $\mathbb{R}_{>0}$ |
| Clipping value of $\tau^2$ | An upper limit on $\tau^2$ | SMDS | $\mathbb{R}_{>0}$ |
| $\kappa$ | GP lengthscale | CLDS | $\mathbb{R}_{>0}$ |
| $\sigma$ | GP scale | CLDS | $\mathbb{R}_{>0}$ |
| $L$ | Number of basis functions for GP approximation | CLDS | $\mathbb{Z}_{\geq 1}$ |

**Table 2:** Experiment-specific Hyperparameter Search.

| Hyperparameter | Sim. Exp. 1<br>(Sec. 4.1) | Sim. Exp. 2<br>(Sec. 4.1) | Macaque Data<br>(Sec. 4.2) | Rodent Data<br>(Sec. 4.3) |
| --- | --- | --- | --- | --- |
| $D$ (LDS) | $\{1, 2, \dots, 10\}$ | $\{4, 6, \dots, 16\}$ | $\{5, 10, \dots, 35\}$ | $\{1, 2, \dots, 10\}$ |
| $D$ (SMDS) | $\{1, 2, 3, 4\}$ | $\{4, 6, \dots, 12\}$ | $\{5, 10, \dots, 35\}$ | $\{1, 2, \dots, 10\}$ |
| $D$ (CLDS) | N/A | N/A | $\{5, 10, \dots, 35\}$ | $\{1, 2, \dots, 10\}$ |
| $\alpha$ | 1e-6 | $\{1\text{e}0, 5\text{e}0, 7\text{e}0, 1\text{e}1, 2\text{e}1, 3\text{e}1, 5\text{e}1\}$ | $\{1\text{e}-3, 1\text{e}-2, 1\text{e}-1\}$ | $\{1\text{e}-6, 1\text{e}-5, 1\text{e}-3\}$ |
| $\beta$ | 1e-6 | 1e-9 | 1e-6 | $\{1\text{e}-5, 1\text{e}-4\}$ |
| Initial $\tau^2$ | 1e-6 | $\{1\text{e}-8, 5\text{e}-8, 1\text{e}-7, 5\text{e}-7, 1\text{e}-6\}$ | $\{5\text{e}-6, 7\text{e}-6, 1\text{e}-5, 2\text{e}-5\}$ | $\{5\text{e}-7, 1\text{e}-6, 5\text{e}-6, 1\text{e}-5\}$ |
| Clipping value of $\tau^2$ | 1e-3 | 1e-4 | 1e-4 | 1e-4 |
| $\kappa$ | N/A | N/A | $\{0.4, 0.45, 0.5\}$ | $\{0.4, 0.45, 0.5\}$ |
| $\sigma$ | N/A | N/A | 0.2 | 0.2 |
| $L$ | N/A | N/A | $\{5, 9, 13, 17, 65, 71, 77, 83\}$ | $\{5, 9, 13, 17, 65, 71, 77, 83\}$ |

### G Simulated Data Experiments

We provide details on our simulated data experiments below. For further information on hyperparameter selection, runtime, and computational resources used in the experiments, please refer to Sec. J.

#### G.1 Toy Experiment

We set the ground truth latent dimensionality to *D* = 2 and the observation dimension to *N* = 10. We set the number of trial conditions to 4. For each condition, we sampled the initial mean of the latent states from a Gaussian distribution with zero mean and diagonal covariance with diagonal values set to 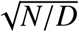. The dynamics matrix was set to a random rotation matrix, and the dynamics and emissions noises were set to 0.01. To generate drift, we sampled displacements 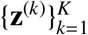 for *K* = 750 trials from a random walk, where the drift rates were sampled independently per dimension: 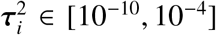, 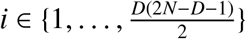. The drift rate τ^2^ for the true model was then set to the empirical variance of the smoothed displacements. A total of 125 trials were left out for test data. We fitted both models for 200 EM iterations. Fig. 6 shows the normalized Grassmann distance matrix computed with the simulated emission matrices (left) and with the learned emission matrices (right). SMDS accurately recovers the ground truth drift.

**Figure 6:**
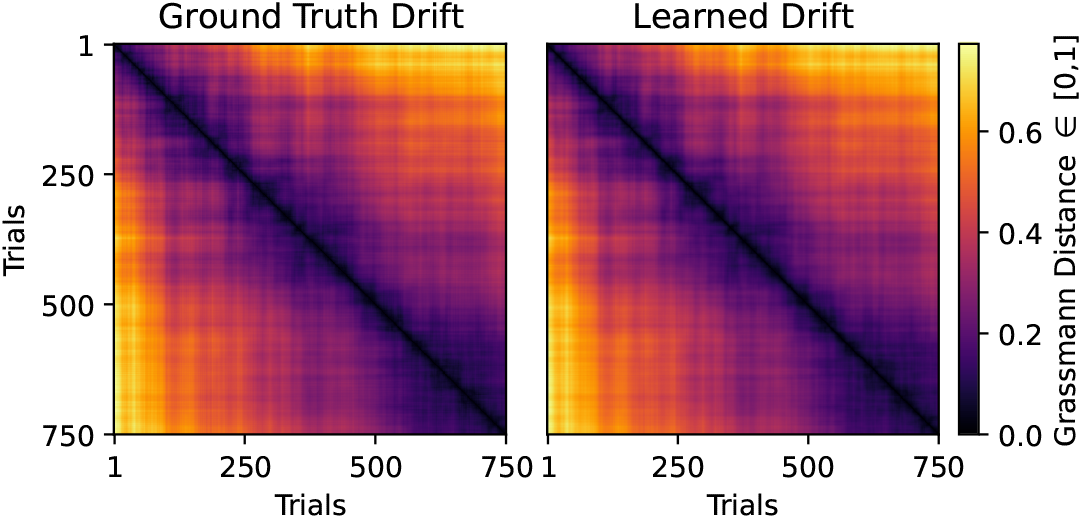
Simulated and learned drift for the toy experiment in Sec. 4.1: We simulated smooth drift and computed the normalized Grassmann distance matrix of the simulated emission matrices (left). We then fitted an SMDS and computed the normalized Grassmann distance matrix with the learned emission matrices. SMDS accurately recovers the simulated drift.

#### G.2 Higher-dimensional Simulated Data Experiment

We validated whether SMDS could recover latent dynamics and observation drift in higher-dimensional setups. We simulated data from an SMDS with *D* = 8 and *N* = 24. We fitted both LDS and SMDS across a range of state dimensions, *D* ∈ [4, 16]. As shown in Fig. 7A, SMDS accurately identified the true latent dimensionality of 8, while we once again observed that the test log-likelihood of LDS continued to increase beyond the true dimension and underperformed relative to SMDS. We also found that SMDS accurately recovered the eigenvalues of the true dynamics matrix (Fig. 7B), whereas LDS failed to do so.

**Figure 7:**
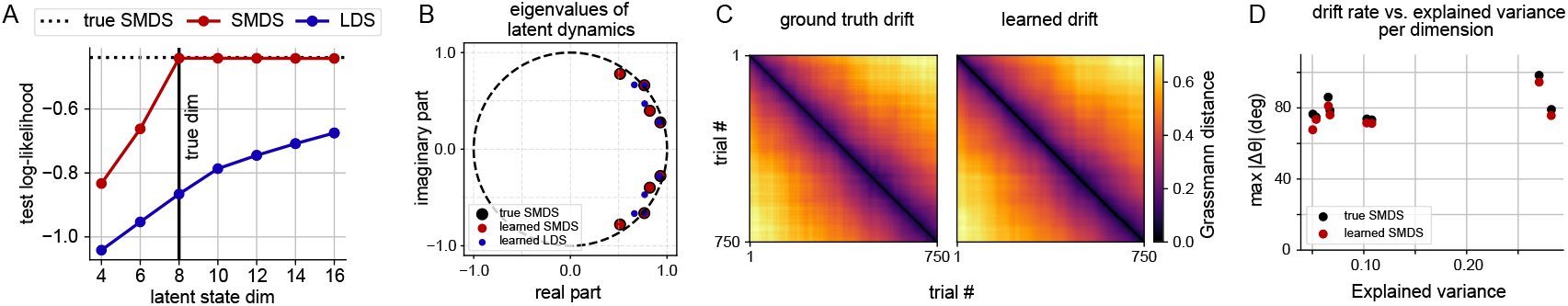
Simulated data experiment 2. **(A)**: Held-out log-likelihood on data simulated from SMDS with *D* = 8, *N* = 24. SMDS outperforms standard LDS and recovers the true state dimension. **(B)**: Eigenvalues of the true dynamics matrix, *A*, compared to those learned by SMDS and LDS. SMDS recovers these accurately. **(C)**: SMDS also recovers ground truth drift, measured by the normalized Grassmann distance across 750 trials. **(D)**: We show the relationship between peak drift and explained variance across individual subspace dimensions from the ground truth data. SMDS accurately recovers this relationship.

**Figure 8:**
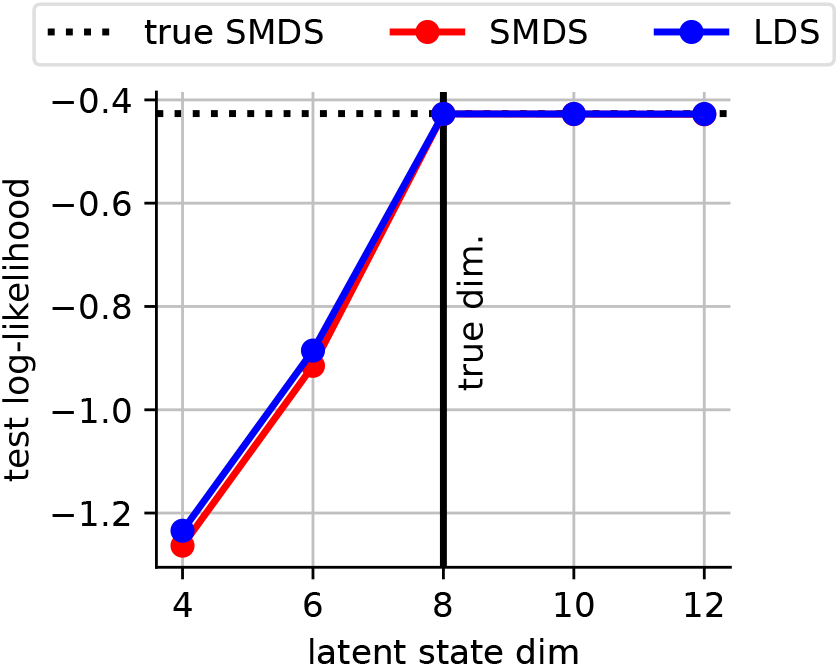
Test log-likelihood plot for the LDS simulated experiment in Sec. 4.1: We simulated data from an LDS with *D* = 8 and *N* = 24. We fitted both LDS and SMDS with state dimensions ranging from 4 to 12.

To assess the ability of SMDS to recover drift, we computed pairwise Grassmann distances (described in Sec. 2.3) using the ground-truth and learned emission matrices between trials (Fig. 7C). We found that SMDS accurately captured the drift pattern in the ground-truth data. We also evaluated whether SMDS could recover the relationship between peak drift (defined in Appendix E) and explained variance across individual subspace dimensions. We made this relationship identifiable by rotating the emission matrices to order the dimensions by their average explained variance. As shown in Fig. 7D, SMDS recovered the ground-truth trend accurately.

These experiments demonstrate the ability of SMDS to accurately capture latent dynamics in the presence of drift. SMDS recovers the correct latent dimensionality and obtains a higher held-out test log-likelihood than an LDS. LDS requires more latent dimensions for the same dataset, highlighting the need to account for drift. SMDS also allows for precise quantification of drift in the observation space.

#### G.3 Experiments with Data Simulated from an LDS

As a control, we simulated stationary data from an LDS (*D* = 8, *N* = 24) and fitted both LDS and SMDS. The emission matrix was set randomly with a standard normal distribution, and the dynamics matrix was set to a random rotation matrix. The dynamics and emissions noises were set to 0.1. We sampled a total of 750 trials, each of which was 30 timesteps long. We fitted SMDS and LDS with state dimensions ranging from 4 to 12. For SMDS, we set the *α* and *β* for the prior on τ^2^ to 1e2 and 1e−9, respectively, and the initial value of τ^2^ to 1e−9. The clipping value of τ^2^ was set to 1e−4. Both models achieved peak test log-likelihood at the true latent dimensionality. LDS slightly outperformed SMDS due to the additional flexibility in modeling displacement dynamics for stationary data. This further validates our inference procedure and demonstrates that SMDS appropriately captures stationary datasets.

### H Modeling Macaque Neural Data

We include further details for the macaque neural data example presented in Sec. 4.2. For further information on hyperparameter selection, runtime, and computational resources used in the experiments, please refer to Sec. J.

#### H.1 Training Details and Hyperparameters

We chunked the trials into blocks of 8 trials, resulting in a total of 93 blocks. Keeping the 8 blocks at each end of the session as training data, we randomly sampled 12 blocks from the remaining blocks for test data across 5 different seeds. To prevent unstable inference caused by overfitting to some channels, we clipped the entries of the diagonal emission covariance matrix to a minimum of either 5e−3 or 4e−3. We fitted both models for 300 EM iterations, and saw that the training log-likelihood saturated.

### I Modeling Rodent Neural Data

We include further details for the rodent neural data example presented in Sec. 4.3. For further information on hyperparameter selection, runtime, and computational resources used in the experiments, please refer to Sec. J.

#### I.1 Training Details and Hyperparameters

We chunked the trials into blocks of 4 trials, resulting in a total of 31 blocks. Keeping the 6 blocks at each end of the session as training data, we randomly sampled 3 blocks from the remaining blocks for test data across 3 different seeds. We fitted both models for 600 EM iterations, and ensured that the training log-likelihood saturated.

#### I.2 PCA Cumulative Explained Variance

We show the PCA cumulative explained variance plot as a function of the number of principal components (Fig. I). We see a kink at around 10 principal components, hence we fit SMDS and LDS with state dimensions from 1 to 10 in our main experiment in Sec. 4.3.

### J Experimental Configurations

Unless otherwise specified, we performed a grid search over a range of values within the permissible set. In certain circumstances, the hyperparameter was selected to consider empirical properties of the data, e.g. we used *D* from 1 to 10 for the rodent neural data based on our observation in Section 9. In addition, for fair comparison, we initialized both **U**_base_ of SMDS and the emission matrix of LDS with PCA. Below is a table that lists the hyperparameter search details for Sec. 4.1, 4.2, and 4.3.

### K PCA Fails to Recover Drift Per Dimension

We set the ground truth latent dimensionality to *D* = 2 and the observation dimension to *N* = 10. The dynamics matrix was set to a random rotation matrix, and the dynamics and emissions noises were set to 0.01. To generate smooth drift, we first sampled displacements 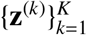 for *K* = 250 trials from a random walk, where the drift rates were sampled independently per dimension: 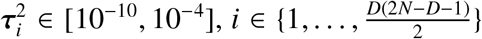. The drift rate τ^2^ for the true model was then set to the empirical variance of the smoothed displacements. Each trial was 200 timesteps long.

As shown in Figure 10, SMDS was able to recover both the overall drift and the drift per individual dimension. On the other hand, PCA accurately captured the overall amount of drift but failed to track how individual dimensions drifted over trials. This shows the failure modes of PCA: the principal components can flip signs or swap their relative ordering between trials, preventing PCA from recovering the true per-dimension drift.

**Figure 9:**
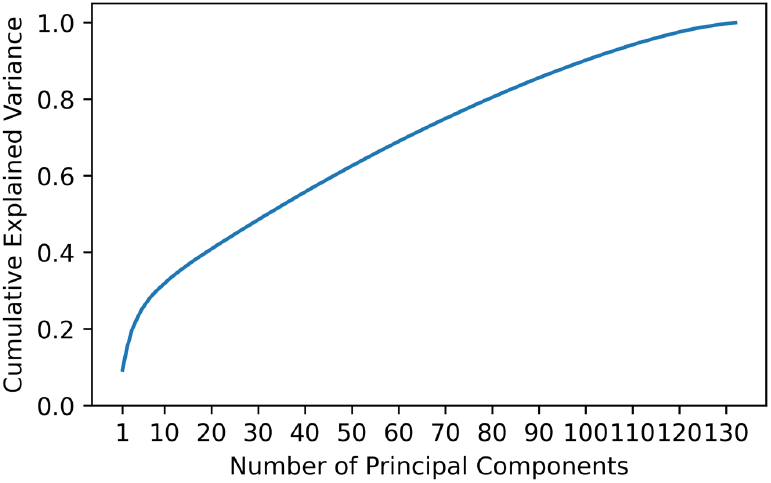
Cumulative explained variance as a function of number of principal components.

**Figure 10:**
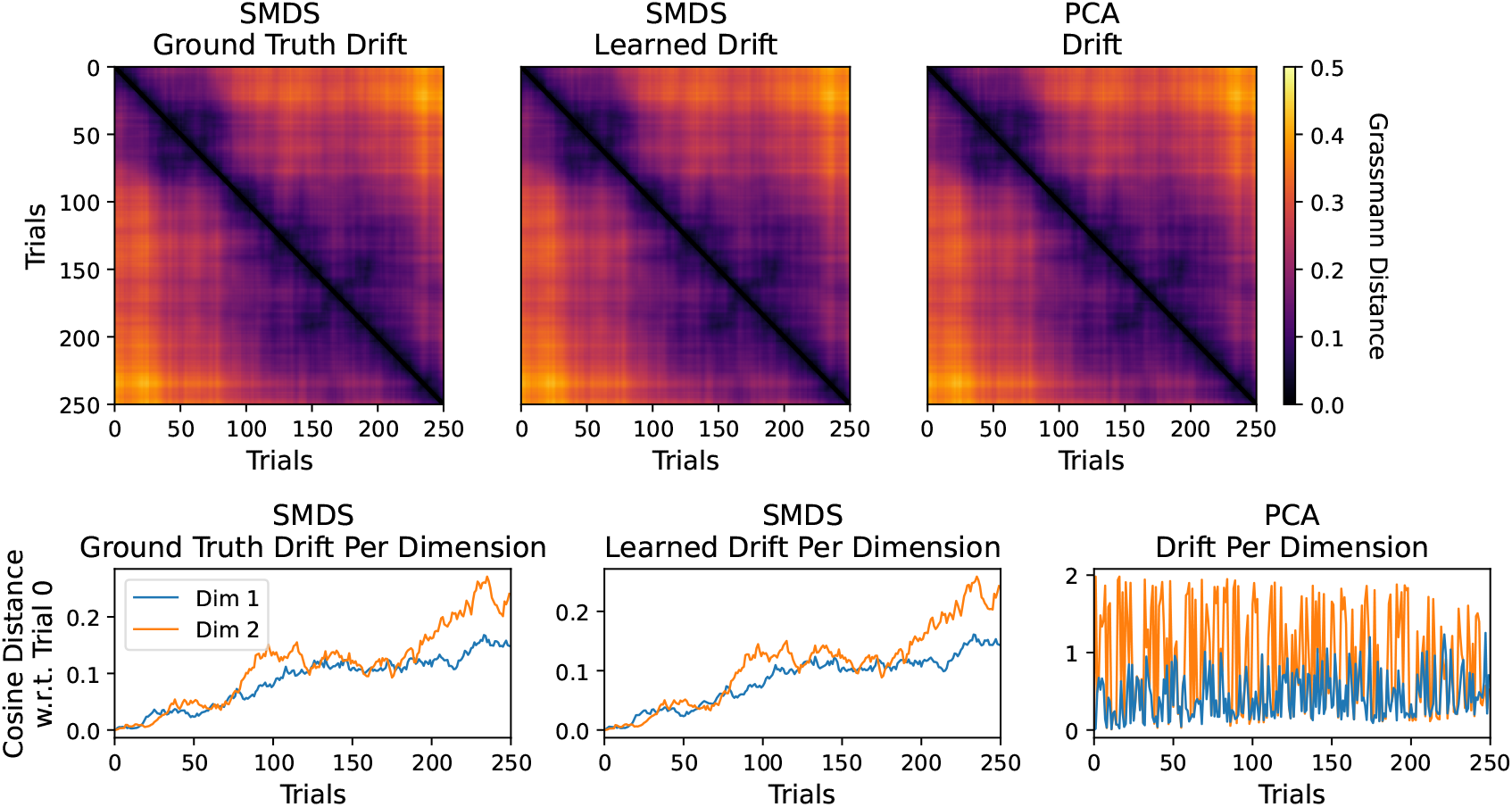
PCA fails to recover per-dimension drift. **(Top Row):** Ground-truth drift from SMDS (left), drift estimated by SMDS (center), and drift estimated by PCA (right). Both SMDS and PCA are able to recover the true overall drift measured by the normalized Grassmann distance. **(Bottom Row):** Drift per dimension from true SMDS (left), learned SMDS (center), and PCA (right). While SMDS accurately recovers the true drift per dimension, measured by cosine distance relative to the first trial, PCA does not.

### L Code Availability

Our implementation of SMDS is available at https://github.com/lindermanlab/smds. The repository includes the source code for SMDS (in Python) and an example Jupyter Notebook. This notebook shows how to sample data from an SMDS and then re-fit an SMDS with the procedure described in Sec. 3.2. SMDS is built based on the framework provided by Dynamax Linderman et al. (2025).

## Footnotes

1 Code for the SMDS model and experiments is available at https://github.com/lindermanlab/smds.

2 Please refer to Clarke et al. (2026) for further details on the dataset and instructions on data access. The dataset ID used in this work is U201202_01.

3 https://github.com/neurostatslab/clds.

## References

Archer, E. W., Koster, U., Pillow, J. W., and Macke, J. H. (2014). Low-dimensional models of neural population activity in sensory cortical circuits. In Advances in Neural Information Processing Systems, volume 27. Curran Associates, Inc.

Balzano, L., Chi, Y., and Lu, Y. M. (2018). Streaming pca and subspace tracking: The missing data case. Proceedings of the IEEE, 106(8):1293–1310.

Bendokat, T., Zimmermann, R., and Absil, P.-A. (2024). A grassmann manifold hand-book: Basic geometry and computational aspects. Advances in Computational Mathematics, 50(1):6.

Bienstock, D., Jeong, M., Shukla, A., and Yun, S.-Y. (2022). Robust streaming pca. Advances in Neural Information Processing Systems, 35:4231–4243.

Björck, Å. and Golub, G. H. (1973). Numerical methods for computing angles between linear subspaces. Mathematics of computation, 27(123):579–594.

Blei, D. M., Kucukelbir, A., and McAuliffe, J. D. (2017). Variational inference: A review for statisticians. J. Am. Stat. Assoc., 112(518):859–877.

Blocker, C. J., Raja, H., Fessler, J. A., and Balzano, L. (2023). Dynamic subspace estimation with grassmannian geodesics. arXiv preprint arXiv:2303.14851.

Bong, H., Liu, Z., Ren, Z., Smith, M., Ventura, V., and Kass, R. E. (2020). Latent dynamic factor analysis of high-dimensional neural recordings. Advances in neural information processing systems, 33:16446–16456.

Bordin, C. J. and Bruno, M. G. (2020). Particle filtering on the complex stiefel manifold with application to subspace tracking. In ICASSP 2020-2020 IEEE International Conference on Acoustics, Speech and Signal Processing (ICASSP), pages 5485–5489. IEEE.

Carmena, J. M., Lebedev, M. A., Henriquez, C. S., and Nicolelis, M. A. (2005). Stable ensemble performance with single-neuron variability during reaching movements in primates. Journal of Neuroscience, 25(46):10712–10716.

Chen, T.-W., Li, N., Daie, K., and Svoboda, K. (2017). A map of anticipatory activity in mouse motor cortex. Neuron, 94(4):866–879.

Chikuse, Y. (2006). State space models on special manifolds. Journal of Multivariate Analysis, 97(6):1284–1294.

Clarke, S. E., Jun, E. J., and Nuyujukian, P. (2026). Neural modes in motor cortex cycle over fast timescales. bioRxiv, pages 2026–01.

Costacurta, J., Duncker, L., Sheffer, B., Gillis, W., Weinreb, C., Markowitz, J., Datta, S. R., Williams, A., and Linderman, S. (2022). Distinguishing discrete and continuous behavioral variability using warped autoregressive hmms. Advances in neural information processing systems, 35:23838–23850.

Dahmen, D., Layer, M., Deutz, L., Dąbrowska, P. A., Voges, N., Von Papen, M., Brochier, T., Riehle, A., Diesmann, M., Grün, S., et al. (2022). Global organization of neuronal activity only requires unstructured local connectivity. elife, 11:e68422.

Driscoll, L. N., Duncker, L., and Harvey, C. D. (2022). Representational drift: Emerging theories for continual learning and experimental future directions. Current Opinion in Neurobiology, 76:102609.

Driscoll, L. N., Pettit, N. L., Minderer, M., Chettih, S. N., and Harvey, C. D. (2017). Dynamic reorganization of neuronal activity patterns in parietal cortex. Cell, 170(5):986–999.e16.

Duncker, L. and Sahani, M. (2018). Temporal alignment and latent gaussian process factor inference in population spike trains. Advances in neural information processing systems, 31.

Duncker, L. and Sahani, M. (2021). Dynamics on the manifold: Identifying computational dynamical activity from neural population recordings. Current Opinion in Neurobiology, 70:163–170.

Eden, U. T., Frank, L. M., Barbieri, R., Solo, V., and Brown, E. N. (2004). Dynamic analysis of neural encoding by point process adaptive filtering. Neural computation, 16(5):971–998.

Gallego, J. A., Perich, M. G., Chowdhury, R. H., Solla, S. A., and Miller, L. E. (2020). Long-term stability of cortical population dynamics underlying consistent behavior. Nature neuroscience, 23(2):260–270.

Gao, Y., Archer, E. W., Paninski, L., and Cunningham, J. P. (2016). Linear dynamical neural population models through nonlinear embeddings. In Advances in Neural Information Processing Systems, volume 29. Curran Associates, Inc.

Geadah, V., Nejatbakhsh, A., Lipshutz, D., Pillow, J. W., and Williams, A. H. (2025). Modeling neural activity with conditionally linear dynamical systems. arXiv preprint arXiv:2502.18347.

Ghahramani, Z. and Hinton, G. E. (2000). Variational learning for switching state-space models. Neural computation, 12(4):831–864.

Glaser, J., Whiteway, M., Cunningham, J. P., Paninski, L., and Linderman, S. (2020). Recurrent switching dynamical systems models for multiple interacting neural populations. In Advances in Neural Information Processing Systems, volume 33, page 14867–14878. Curran Associates, Inc.

Hu, A., Zoltowski, D., Nair, A., Anderson, D., Duncker, L., and Linderman, S. (2024). Modeling latent neural dynamics with gaussian process switching linear dynamical systems. Advances in Neural Information Processing Systems, 37:33805–33835.

Jha, A., Gupta, D., Brody, C., and Pillow, J. W. (2024). Disentangling the roles of distinct cell classes with cell-type dynamical systems. Advances in Neural Information Processing Systems, 37:33668–33690.

Lee, H. D., Warrington, A., Glaser, J., and Linderman, S. (2023). Switching autoregressive low-rank tensor models. Advances in Neural Information Processing Systems, 36:57976–58010.

Linderman, S., Johnson, M., Miller, A., Adams, R., Blei, D., and Paninski, L. (2017). Bayesian learning and inference in recurrent switching linear dynamical systems. In Artificial intelligence and statistics, pages 914–922. PMLR.

Linderman, S. W., Chang, P., Harper-Donnelly, G., Kara, A., Li, X., Duran-Martin, G., and Murphy, K. (2025). Dynamax: A python package for probabilistic state space modeling with jax. Journal of Open Source Software, 10(108):7069.

Macke, J. H., Buesing, L., Cunningham, J. P., Yu, B. M., Shenoy, K. V., and Sahani, M. (2011). Empirical models of spiking in neural populations. In Advances in Neural Information Processing Systems, volume 24. Curran Associates, Inc.

Marks, T. D. and Goard, M. J. (2021). Stimulus-dependent representational drift in primary visual cortex. Nature Communications, 12(1):5169.

Mudrik, N., Chen, Y., Yezerets, E., Rozell, C. J., and Charles, A. S. (2024). Decomposed linear dynamical systems (dlds) for learning the latent components of neural dynamics. Journal of Machine Learning Research, 25(59):1–44.

Narayanamurthy, P. and Vaswani, N. (2018a). Nearly optimal robust subspace tracking. In International Conference on Machine Learning, pages 3701–3709. PMLR.

Narayanamurthy, P. and Vaswani, N. (2018b). Provable dynamic robust pca or robust subspace tracking. IEEE Transactions on Information Theory, 65(3):1547–1577.

Nejatbakhsh, A., Garon, I., and Williams, A. (2023). Estimating noise correlations across continuous conditions with wishart processes. Advances in Neural Information Processing Systems, 36:54032–54045.

Oja, E. (1982). Simplified neuron model as a principal component analyzer. Journal of mathematical biology, 15:267–273.

Pandarinath, C., O’Shea, D. J., Collins, J., Jozefowicz, R., Stavisky, S. D., Kao, J. C., Trautmann, E. M., Kaufman, M. T., Ryu, S. I., Hochberg, L. R., et al. (2018). Inferring single-trial neural population dynamics using sequential auto-encoders. Nature methods, 15(10):805–815.

Paninski, L., Ahmadian, Y., Ferreira, D. G., Koyama, S., Rahnama Rad, K., Vidne, M., Vogelstein, J., and Wu, W. (2010). A new look at state-space models for neural data. Journal of computational neuroscience, 29(1):107–126.

Rentmeesters, Q., Absil, P.-A., Van Dooren, P., Gallivan, K., and Srivastava, A. (2010). An efficient particle filtering technique on the grassmann manifold. In 2010 IEEE International Conference on Acoustics, Speech and Signal Processing, pages 3838–3841. IEEE.

Rubin, A., Geva, N., Sheintuch, L., and Ziv, Y. (2015). Hippocampal ensemble dynamics timestamp events in long-term memory. eLife, 4:e12247.

Rule, M. E., Loback, A. R., Raman, D. V., Driscoll, L. N., Harvey, C. D., and O’Leary, T. (2020). Stable task information from an unstable neural population. eLife, 9:e51121.

Rule, M. E., O’Leary, T., and Harvey, C. D. (2019). Causes and consequences of representational drift. Current Opinion in Neurobiology, 58:141–147.

Sasfi, A., Padoan, A., Markovsky, I., and Dörfler, F. (2024). Subspace tracking for online system identification. arXiv preprint arXiv:2412.09052.

Smith, A. C. and Brown, E. N. (2003). Estimating a state-space model from point process observations. Neural computation, 15(5):965–991.

Soldado-Magraner, J., Mante, V., and Sahani, M. (2024). Inferring context-dependent computations through linear approximations of prefrontal cortex dynamics. Science Advances, 10(51):eadl4743.

Srivastava, A. and Klassen, E. (2004). Bayesian and geometric subspace tracking. Advances in Applied Probability, 36(1):43–56.

Särkkä, S. (2013). Bayesian Filtering and Smoothing. Institute of Mathematical Statistics Textbooks. Cambridge University Press.

Tompkins, F. and Wolfe, P. J. (2007). Bayesian filtering on the stiefel manifold. In 2007 2nd IEEE International Workshop on Computational Advances in Multi-Sensor Adaptive Processing, pages 261–264. IEEE.

Vyas, S., Golub, M. D., Sussillo, D., and Shenoy, K. V. (2020). Computation through neural population dynamics. Annual Review of Neuroscience, 43(Volume 43, 2020):249–275.

Wen, Z. and Yin, W. (2013). A feasible method for optimization with orthogonality constraints. Math. Program., 142(1–2):397–434.

Yang, B. (1995). Projection approximation subspace tracking. IEEE Transactions on Signal processing, 43(1):95–107.

Zhao, Y. and Park, I. M. (2017). Variational latent gaussian process for recovering single-trial dynamics from population spike trains. Neural computation, 29(5):1293–1316.

Ziv, Y., Burns, L. D., Cocker, E. D., Hamel, E. O., Ghosh, K. K., Kitch, L. J., Gamal, A. E., and Schnitzer, M. J. (2013). Long-term dynamics of ca1 hippocampal place codes. Nature Neuroscience, 16(3):264–266.

Zoltowski, D. M., Pillow, J. W., and Linderman, S. W. (2020). A general recurrent state space framework for modeling neural dynamics during decision-making. In Iii, H. D. and Singh, A., editors, Proceedings of the 37th International Conference on Machine Learning, volume 119 of Proceedings of Machine Learning Research, pages 11680–11691, Virtual. PMLR.

